# Population differences in how wild Trinidadian guppies use social information and socially learn

**DOI:** 10.1101/786772

**Authors:** Laura Chouinard-Thuly, Simon M. Reader

**Author notes:** Corresponding author: Laura Chouinard-Thuly, Department of Biology, McGill University, 1205 Dr. Penfield, Montréal, Québec, Canada, H3A 1B1.

## Abstract

Animals have access to information produced by the behaviour of other individuals, which they may use (“social information use”) and learn from (“social learning”). The benefits of using such information differ with socio-ecological conditions. Thus, population differences in social information use and social learning should occur. We tested this hypothesis with a comparative study across five wild populations of Trinidadian guppies (*Poecilia reticulata*) known to differ in their ecology and social behaviour. Using a field experiment, we found population differences in how guppies used and learned from social information, with only fish from one of the three rivers studied showing evidence of social information use and social learning. Within this river, populations differed in how they employed social information: fish from a high-predation regime where guppies exhibit high shoaling propensities chose the same foraging location than conspecifics, while fish from a low-predation regime with reduced shoaling propensities chose and learned the opposite foraging location than conspecifics. We speculate that these differences are due to differences in predation risk and conspecific competition, possibly mediated via changes in grouping tendencies. Our results provide evidence that social information use and social learning can differ across animal populations and are influenced by socio-ecological factors.

---

Individuals that possess reliable information about resources and threats can make strategic decisions [1]. Animals can gather information from different sources, either by interacting directly with the environment and thus acquiring ‘asocial’ information or from the behaviour or products of other individuals, a process termed social information use. Animals may retain information for later use, and thus learn from personally acquired information or socially acquired information (i.e. social learning [2]). Which type of information to use in a given context is strategic and based on trade-offs [3–5]. The respective costs and benefits depend largely on recent and current socio-ecological conditions such as predation and/or stress level [6,7] or social group composition [8]. It also appears that flexibility in both social information use and social learning can be constrained by individual characteristics [9–11] and shaped by recent experience of the reliability of information [e.g. 12]. Thus, the decision to rely on social sources of information is not be solely dependent on reliability or net benefit of the information in the current situation, but also by individual tendencies. Whether these processes translate to differences in social information use between populations dwelling in different socio-ecological environments is rarely investigated, but likely.

Current local conditions shape the costs and benefits of asocial and social information. Using social cues can reduce the energy required to acquire information and is particularly beneficial if energy is limited [13]. Social information reduces risk related to personally sampling a resource [6,7], particularly in a context with predation pressure. In other cases, using social cues can be maladaptive or suboptimal if the information gathered is outdated or irrelevant to the observer [14,16], and using social information may increase competition if individuals thus converge on a limited resource [17]. Local current environmental characteristics shape which type of information is most likely to be beneficial [3] and influences the decision of individuals [11,18,19]. If individuals are completely flexible in their decision, current local conditions determine which information individuals should rely on.

However, we find that individuals are constrained in their decision through early-life experience and evolutionary history with the benefits of social information. For example, bumblebees *(Bombus terrestris)* will learn to copy or avoid other individuals’ foraging choices depending on whether following these social cues was previously rewarded, demonstrating the effect of recent experience [20]. Early-life experiences can also shape adult social information use, either due to direct experience with the value of following social cues [e.g. 12], or to broader differences in social experience such as maternal care [21,22]. Species differences in social information use and social learning have also been described (e.g. birds: [23]; mammals: [24]) which could be the result of evolved and/or developmental influences. For example, ninespine sticklebacks (*Pungitius pungitius)* who are under high predation pressure display increased propensities to socially learn than the less predated but closely related threespine sticklebacks (*Gasterosteus aculeatus)* [25,26]. Furthermore, individual behavioural phenotypes that themselves could be shaped by experience and evolution, such as the speed to explore a novel environment or to solve a novel problem, can also predict social information use [10,18,27,28].

Given the short-term and long-term influences on trade-offs between both types of information, differences between populations dwelling in different socio-ecological environments are likely. However, very little work has investigated such population differences, particularly with experimental tests. A notable exception is the finding that populations of Zenaida doves *(Zenaida aurita)* differ in how they learn from a Carib grackle (*Quiscalus lugubris)*, a finding that has been explained by differences in foraging ecology shaping differences in social behaviour between these populations [29,30]. Here, we investigated population differences in social information use and social learning by comparing multiple replicate populations tested in the wild, with the aim of identifying ecological factors that shape social information use.

We used wild Trinidadian guppies to investigate this question. Guppies have successfully colonizing rivers that are extremely diverse in geography and ecology [31]. Guppies readily learn from conspecifics and hetereospecifics in both the field and the laboratory, which may partially explain why they thrive in diverse and new conditions [32,33]. The ecology and evolution of Trinidadian guppies is well studied, with differences in physiology, morphology, life history and behaviour found between populations that are partially separated by natural barriers, driven mostly, but not only, by the presence, density, and composition of predators [34–36]. Upper river habitats in northern Trinidad typically contain fewer predators of adult guppies, as well as a weaker current, and more access to invertebrates than lower river habitats [37]. Trinidadian guppy populations differ on numerous behavioural measures: guppies from the upper river populations display lower shoaling tendencies, higher intraspecific aggressiveness and competition, and bolder phenotypes than in the lower river [38–42]. High shoaling tendencies could increase the propensity to rely on social information since individuals are near conspecifics, while high aggression and competition may increase the net costs of social information use and social learning. Trinidadian populations provide a valuable opportunity to test natural variation in the transmission of social information between populations exposed to varying environments.

We compared propensities for social information use and social learning using a foraging task in five populations of wild Trinidadian guppies from three rivers. Our design allowed us to investigate not only if there are population differences, but also whether within-river differences were paralleled across different rivers, which would provide support for socio-ecological conditions shaping population differences in a consistent manner. We predicted guppies to prefer to forage at the same location as conspecific demonstrators, and to retain this preference when demonstrators were removed, as previously shown [33]. However, we expected these tendencies to vary across populations. In fish from the Lower Aripo, known to display high shoaling tendencies and low interspecific aggression [37], and from the Lower Marianne, we predicted subjects would copy the demonstrated location. In comparison, we expected guppies from the Upper Aripo, Upper Marianne and Paria, known to display low shoaling tendencies and expected (Marianne) or shown (Upper Aripo, Paria) to show high interspecific aggression [37], to either avoid the demonstrated location or to be unaffected by social cues. Guppies from the Paria site show particularly low shoaling tendencies and high interspecific aggression, making it an interesting comparator [42]. We expected similar population differences between the Upper and Lower sites in the Aripo and Marianne rivers, although recent literature suggests that rivers may not be perfect replicates [43]. This comparative study of social information use and social learning propensities thus allows us to determine (1) whether populations differ in these propensities, as might be predicted from hypotheses that evolutionary and developmental processes shape social information use; (2) why and when propensities change, and (3) whether these propensities change in similar manner, thus providing evidence for specific socio-ecological factors shaping social information use.

## Methods

### Overview

We used a foraging test to compare how five guppy populations used social information and learned from conspecifics. We assessed social information use and social learning by (1) comparing subjects’ responses to conspecific ‘demonstrators’ at two feeding locations in a counterbalanced design and (2) comparing these responses to control subjects not exposed to demonstrators. Social information use was measured during a demonstration phase, when demonstrators were present (except in the control trials), while social learning was measured during a subsequent test phase, when demonstrators had been removed. Social influences on behaviour would result in subjects being more or less likely to feed at the demonstrated location than the alternative location.

### Study sites and sampling

We tested in three rivers located in different watersheds of the Northern Range Mountains in Trinidad: the South slope Aripo river (June 2013), the North slope Paria river (June 2013), and the North slope Marianne river (July 2014). We tested at previously studied sites (Ar2 Ar4, Ma14, Ma8, Pa14) detailed in [44] and [45]. Guppy lineages from these rivers are genetically differentiated [46]. ‘Upper’ and ‘Lower’ river locations from the Aripo and Marianne rivers are separated by waterfalls, with large teleost fish predators absent from upper but not lower locations, and numerous other ecological differences between the locations [37]. There is no similar ‘Lower’ location in Paria, so we thus sampled only one site that has no large teleost fish predators (similar to other ‘Upper’ locations), but where large predatory prawns *Macrobrachium crenulatum* are present [47,48]. We chose sites where the ectoparasite *Gyrodactylus* has been recorded [44,45].

To ensure independent fish were sampled, we typically selected subsequent sampling pools by going upstream, or by selecting physically separated pools. We used butterfly nets to gently collect female guppies, and ran our tests in enclosures within rivers. Fish were held in a water-filled enclosure placed in the river for a maximum of 5 hours. During this time, we presented them with the social information use and learning tests, then moved them to an enclosure for tested fish, with fish released at their capture site at the end of a testing day.

### Testing apparatus

The testing apparatus consisted of a small floating box made of mosquito net (23 cm high, 38 cm wide and long), which allowed stream water to flow freely through the apparatus, with the front and back of the apparatus made of transparent plastic. Since fish were tested in an enclosure, they were physically separated from any local predators and the experiment was not a field test of social learning on free-living animals [49]. However, they were in field conditions until the experiment began, were tested in their local environment, and were exposed to olfactory and visual cues from outside the enclosure. We mounted a waterproof camera (1080p at 30fps, GoPro3 Black Edition, San Mateo, California) on one wall to record behaviour at the removable feeder (36 cm width) positioned on the opposite side. This feeder consisted of two feeding locations separated by 10 cm, with each location made up of two vertical 5 cm wide feeding columns placed 3 cm apart, creating patches of food that were accessible to multiple individuals simultaneously. The feeding columns were made of food sprinkled on gelatin (KNOX, Treehouse Foods, New York State, USA) mixed with food colouring (Club House, McCormik Canada, London Ontario, Canada) poured on a patterned background. We created two types of feeding column on the feeding wall. One was made of freeze-dried bloodworms (*Chironomus* spp., Omega One, Omegasea Ltd, Sitka, Alaska) sprinkled on green-coloured gelatin, placed on a black-striped background. The other was made of flake food (TetraMin, Tetra, Germany) sprinkled on yellow-coloured gelatin placed on a black-dotted background. We used a variety of food, pattern, location and colour cues to provide multiple discriminatory cues for the subjects and to increase differences between the feeding columns. For demonstrations by conspecifics, we put “demonstrator” fish in a small “demonstration box” (10 cm height, 5 cm width and depth) made of perforated transparent plastic so that demonstrators reliably fed on one column without requiring extensive training. We placed the box directly in front of one column, with a similar but smaller feeding column inside the box.

### Experimental methods

Each trial consisted of a 1) habituation, 2) demonstration, and 3) test phase. In the 1) habituation phase, we placed a group of four fish in the testing apparatus without the feeding wall for 10 minutes. We tested fish in groups as guppies are typically highly social and may show population dependant stress responses when placed in isolation [41,50], potentially impacting the social information use we examine here. Simultaneously, two fish from the previously tested subject group, selected at random to act as demonstrators, were habituated to the demonstration box outside of the apparatus. All demonstrators fed during this phase. Between the habituation phase and the demonstration phase, we inserted an opaque partition between the fish and the foraging area. With the partition in place, we inserted the feeder wall and demonstration box out of view of the subjects. The demonstration box was placed in front of one of the four columns, and thus at one of the two locations and at one of the two column types, except for the control groups which viewed no demonstrators. The control groups were run twice per testing day, as the first test each day (thus providing demonstrators for the first demonstration of the day) and a second test chosen at random. We counterbalanced the demonstration groups between the four columns every day. The 2) demonstration phase started upon lifting of the partition and lasted 6 minutes and was used to determine the propensity of subjects to use social information. During this phase we allowed fish to freely move and access the food resources. This procedure differs from many social learning tests where subjects only observe feeding behaviour (but see [49] for similar procedures). We considered it important to maintain ecological relevance and match much guppy foraging in the wild. Moreover, blocking subject access to food could represent a situation where conspecifics prevent foraging access. Between the demonstration and test phase, the opaque partition was reinserted, the feeding sheet rinsed to remove any odour cues and placed inverted (to reverse the order of the columns and further remove odour biases), and the demonstration box was removed. The 3) test phase started upon lifting of the partition and lasted 8 minutes, and was used to evaluate if social learning had occurred. As on the demonstration phase, the subjects could feed and were rewarded at any foraging location.

From the video recordings, one of two observers blind to the population tested counted the number of feeding pecks [51] on each food column. Since we could not discriminate individuals, we summed the feeding pecks of the four subjects tested together as a group. No feeding pecks were observed away from the food columns. Inter-observer reliability was measured for 30 videos and was high (ICC= 0.81, 95% C.I. = 0.73 < ICC < 0.86). In total, we tested 82 groups with demonstrators and 25 control groups. Of these, 17 were from the Lower Aripo, 15 from the Upper Aripo, 33 from the Lower Marianne, 30 from the Upper Marianne, and 12 from the Paria.

### Statistical analyses

All statistical analyses were performed using R version 3.2.2 [52] and the packages ggplot2 [53] and lme4 [54]. We found no evidence that the demonstrated feeding column type affected foraging behaviour (unpublished data), and thus below we examined feeding locations and feeding rate only.

#### Population differences

We wanted to investigate if and why populations differed in social information use and social learning. We thus examined the influence of demonstrator location on subjects’ foraging location choices for the fish exposed to demonstrators. We ran generalized linear mixed-effect models (GLMMs) with a binomial distribution with the distribution of pecks between the demonstrated and the undemonstrated location as the response variable for the demonstration phase, and for the test phase. This approach, compared to examining the total or percentage of pecks at the demonstrated location, accounts for differences in groups’ propensities to feed.

We investigated population differences between the five sites we tested: Lower Aripo, Upper Aripo, Lower Marianne, Upper Marianne, and Paria. The model also included an observation-level random effect to correct for overdispersion [55]. In this model, an effect of site indicates that populations differed in in their proportion of pecks at the demonstrated location. The reference for the site model was the Lower Aripo, so this site model already provided a population comparison within the river Aripo. We followed by specifically investigating population differences within the Marianne river, by running a GLMM that included the main effect ‘population’ (‘Upper’ or ‘Lower’). We only had one population in the river Paria, so we did not do any follow-up analysis.

#### Social information use and social learning

While the site model above examined whether populations differed in their reaction to the demonstrators, we also need to know how they reacted to the demonstrator. If demonstrator location had no influence, we would expect subjects to peck equally at both locations. We therefore tested whether the observed distribution of pecks differed from chance expectation, which we set at 50% assuming fish randomly feed at both feeders. We did this by removing the intercept of the site model, thus forcing the model to compare the population’s estimates to zero on the latent scale or 50% on the original scale.

#### Feeding rate

To investigate whether demonstrator presence changed the total number of pecks subjects performed (i.e. feeding rate), we ran generalised linear mixed effect model (GLMM) with a Poisson distribution for each river and each phase. Rivers rather than populations were analysed so that an adequate amount of control data was available. The models had the response variable ‘total pecks’, and the main effect ‘demonstration’ (“control” or “with demonstration”) to compare the absolute number of pecks of fish from the control group to the fish with a demonstration. We included as random effects population and group as well as an observation-level random effect to correct for overdispersion. A significant main effect of demonstration with a positive estimate would indicate that exposure to demonstrators increased feeding rate.

#### Feeding location consistency

To analyse whether control fish acquired a preference about feeding locations regardless of social cues, we analysed whether the group random effect significantly helped explain a significant part of the variation. We did this by creating an overall river model for the control trials. In the model, we included the main effect ‘river’ and ‘phase’ to create a repeated measure model. Using a likelihood ratio test (LRT), we compared the overall river model with the same model from which we removed the group random effect, to evaluate if a significant amount of variation is explained by groups.

## Results

### Site differences

#### Demonstration phase

During the demonstration phase, which examined differences in social information use, sites varied in the proportion of pecks at the demonstrated location: fish from the Upper Aripo, Lower Marianne and Paria pecked significantly less (*P* = 0.0046; *P* = 0.028; *P* = 0.018, respectively; table 2; figure 1) at the demonstrated location than our reference site Lower Aripo. Examining the two Marianne populations alone, there were no significant differences in the proportion of pecks at the demonstrated location during the demonstration phase (table 2).

**Table 1:** Site differences in proportion of pecks at the demonstrated location. Estimates and standard error of fixed parameters and their interaction for the GLMM looking at the effect of site on the proportion of pecks at the demonstrated location, defined as a binomial variable of number of ‘successes’ (proportion pecks at demonstrated location) and number of ‘misses’ (proportion pecks at the undemonstrated location). Estimates are presented on the logit scale. The reference level was Lower Aripo for “site”. The model also included an observation-level random effect to correct for overdispersion.

|  | Demonstration phase |  |  |  | Test phase |  |  |  |
| --- | --- | --- | --- | --- | --- | --- | --- | --- |
|  | Estimate | Std. Error | z value | P-value | Estimate | Std. Error | z value | P-value |
| Intercept | 3.64 | 1.70 | 2.14 | **0.033** | 1.81 | 1.23 | 1.47 | 0.14 |
| Upper Aripo | -6.07 | 2.14 | 2.83 | **0.0046** | -4.16 | 1.60 | 2.60 | **0.0093** |
| Paria | -5.53 | 2.33 | 2.38 | **0.018** | -2.13 | 1.70 | 1.26 | 0.21 |
| Lower Marianne | -4.41 | 2.01 | 2.20 | **0.028** | -0.75 | 1.50 | 0.50 | 0.62 |
| Upper Marianne | -3.33 | 2.01 | 1.66 | 0.098 | -1.35 | 1.50 | 0.90 | 0.37 |

**Table 2:** Population differences in the Marianne river. Estimates and standard error of fixed parameters and their interaction for the GLMM looking at the effect of population on the proportion of pecks at the demonstrated location in the Marianne river, defined as a binomial variable of number of ‘successes’ (proportion pecks at demonstrated location) and number of ‘misses’ (proportion pecks at the undemonstrated location). Estimates are presented on the logit scale. The reference level was Lower Marianne for “population”. The model also included an observation-level random effect to correct for overdispersion.

|  | Demonstration phase |  |  |  | Test phase |  |  |  |
| --- | --- | --- | --- | --- | --- | --- | --- | --- |
|  | Estimate | Std. Error | z value | P-value | Estimate | Std. Error | z value | P-value |
| Intercept | -0.70 | 0.85 | 0.82 | 0.41 | 0.94 | 0.69 | 1.35 | 0.18 |
| Population (Upper Marianne) | 1.01 | 1.23 | 0.82 | 0.41 | -0.56 | 0.99 | 0.56 | 0.58 |

**Table 3:** Effect of having a demonstration on total number of pecks of fish. The estimates are presented on the log scale for the demonstration phase (left) and the test phase (right) for the river Aripo (top), Marianne (middle) and Paria (bottom). Our reference levels were no demonstration for the demonstration factorThe GLMM included also a correction for overdispersion in the random effects. Significant p-values (P<0.05) are presented in bold.

|  | Demonstration phase |  |  |  | Test phase |  |  |  |
| --- | --- | --- | --- | --- | --- | --- | --- | --- |
|  | Estimate | Std. Error | z value | P-value | Estimate | Std. Error | z value | P-value |
| **Aripo** |  |  |  |  |  |  |  |  |
| Intercept | -0.71 | 1.09 | 0.65 | 0.52 | 0.63 | 1.05 | 0.60 | 0.55 |
| Demonstration (demonstration) | 2.24 | 1.16 | 1.92 | 0.055 | 1.74 | 1.15 | 1.51 | 0.13 |
| **Marianne** |  |  |  |  |  |  |  |  |
| Intercept | 0.28 | 1.03 | 0.27 | 0.79 | 0.057 | 1.03 | 0.056 | 0.96 |
| Demonstration (demonstration) | -0.22 | 1.14 | 0.20 | 0.84 | 0.61 | 1.10 | 0.55 | 0.58 |
| **Paria** |  |  |  |  |  |  |  |  |
| Intercept | 0.079 | 1.70 | 0.047 | 0.96 | 2.20 | 1.53 | 1.44 | 0.15 |
| Demonstration (demonstration) | 1.64 | 1.82 | 0.90 | 0.37 | 0.041 | 1.73 | 0.024 | 0.98 |

**Figure 1:**
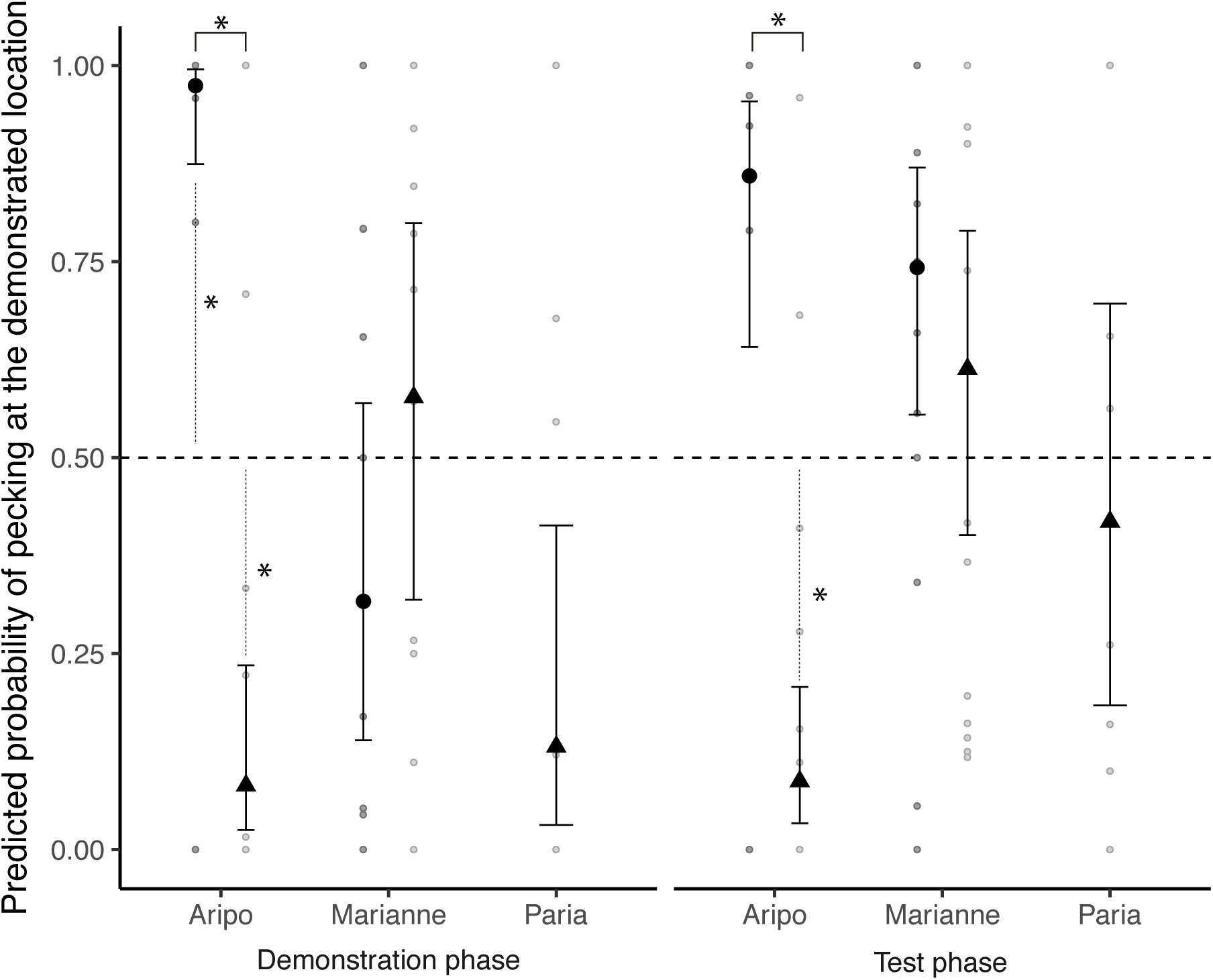
Predicted probability and raw data of pecks (estimate +/- SE) performed at the demonstrated feeder by five populations of fish in the three rivers (circles: lower river; triangle: upper river) for the demonstration and test phases. The dashed line at zero represents our chance expectation of 50% on the original scale. A difference from chance (50%) is indicated by a star and a dotted line, while a difference between populations is indicated by an * above the compared groups. Populations from the Aripo river differed from each other and from chance.

#### Test phase

During the test phase, which examined differences in social learning, Upper Aripo fish pecked significantly less at the demonstrated location compared to Lower Aripo fish (*P* = 0.0093; table 2, figure 1). Other sites did not differ in the proportion of pecks at the demonstrated feeder during the test phase (*Ps* > 0.2; table 2). Examining the two Marianne populations alone, there were no significant differences in the proportion of pecks at the demonstrated location during the test phase (table 2).

### Social information use and social learning

#### Demonstration phase

During the demonstration phase, when demonstrators were present, fish from the Lower Aripo pecked significantly more than expected by chance at the demonstrated location, with 97% of pecks (P = 0.033; figure 1.2, table S1). In contrast, fish from the Upper Aripo pecked significantly less than expected by chance at the demonstrated location, with only 8% of pecks (P = 0.05; table S1). Fish from the Lower Marianne, Upper Marianne, and Paria did not peck at the demonstrated location significantly more or less than the chance expectation of 50%.

#### Test phase

During the test phase, when demonstrators had been removed, fish from the Lower Aripo made 86% of pecks at the previously demonstrated location, but this was not significantly different from chance (*P* = 0.14; figure 1; table S1). Upper Aripo fish made only 9% of pecks at the previously demonstrated location, significantly different from chance (*P* = 0.020; table S1). Fish from the Lower Marianne, Upper Marianne, and Paria did not peck at the previously demonstrated location significantly more or less than the chance expectation.

### Feeding rate

#### Demonstration phase

In the Aripo river, exposure to demonstrators increased slightly the total number of feeding pecks compared to the control condition, but this was not significant (*P* = 0.055; Figure 1). Exposure to demonstrators did not significantly increase the total number of feeding pecks in the other rivers (*P*s > 0.3; Figure 1).

#### Test phase

Exposure to demonstrators did not significantly increase the total number of feeding pecks in any river (*P*s > 0.1; Figure 1).

### Feeding location consistency

There was no evidence that fish from the control groups, without a demonstration, had a consistent preference for a feeding location over the two experimental phases. That is, the model with a group random effect that accounted for repeated measures was not a significantly better fit than the model without for control groups (LRT *X*^2^ = 1.17, df=1, *P* = 0.28). We did find evidence that groups with demonstration had a consistent preference for a feeding location. The model with the group random effect was significantly better at explaining variation than the model without for groups with a demonstration (LRT *X*^2^ = 7.45, df = 1, *P* = 0.006). In other words, only fish with a demonstration showed a consistent preference for a certain feeder.

## Discussion

Using a comparative experiment in wild habitats, we compared the effect of a social demonstration on foraging rate and foraging location across guppy populations. We found that the response to social information varied between populations. We only found evidence for social information use and social learning in fish in the Aripo river. Moreover, within the Aripo river, populations differed in how they reacted to social information: fish from the Upper Aripo avoided the location where conspecifics were seen feeding and retained this bias after the removal of the demonstrators, while fish from the Lower Aripo foraged at the demonstrated location, but this bias was not statistically significant (although still substantial) when demonstrators were removed. Our results show population variation in social information use and social learning, suggesting that evolutionary and/or developmental experiences shape social information use and social learning propensities.

Perhaps our most interesting finding is that Aripo populations reacted differently to social information. Their habitats differ on multiple characteristics, such as food productivity, light levels, and predation pressure, providing multiple possible explanations for the differences we observed. However, predation pressure and competition provide the most likely explanations. The Lower Aripo population is characterised by very cohesive and large shoals, a result of the local predation regime, with little intraspecific aggression [37,42]. In contrast, in the Upper Aripo, predators of adult guppies are mostly absent, and food is more scarce than in the lower reaches [43], with fish displaying lower shoaling tendencies and higher aggression [41]. Thus fish in the Upper Aripo will suffer intraspecific competition if foraging in a group, will gain little in terms of anti-predator benefits, and resource patches may be more rapidly depleted, potentially explaining their tendencies to avoid locations where conspecifics are or were foraging [56]. While most work on social information use has focused on animals matching demonstrator behaviour, animals can employ social information in a variety of ways, including avoiding the choices of others [20,57,58]. Fish in the Lower Aripo suffer increased risks of individual exploration and leaving the group, suffer little intraspecific competition when foraging where others forage, and have easy access to social information, potentially explaining their copying behaviour. Previous work has linked between-individual variation in shoaling tendency with social information use in fish [8,28], and with sociality more broadly in corvids [23]. Competition and limited resources availability has been proposed as an important influence on social information use in species as varied as Japanese quail (*Coturnix japonica*) [57] and fruitfly *Drosophila melanogaster* larvae [59].

Fish from the Aripo, but not the Marianne or Paria rivers, showed evidence for social information use and social learning. Thus, we did not find evidence of parallelism between rivers in this foraging test. A parallel response would have been indicative of a strong effect of specific socio-ecological factors like the presence of predators. Recently, work has highlighted important differences between rivers and drainages in the flow, productivity, and canopy cover [60]. So while certain traits, like coloration, may be selected independently of the composition of the predator community [47,48], some are particular responses to the type, composition, and density of predators [36,43,60,61]. Additionally, other habitat characteristics and how they interplay with predation may be important. For example, guppy density strongly impacts competition and mate choice [62], and light spectrum affects mating tactics [63]. Environmental characteristics that shape competition are particularly likely to shape social information use [56].

Much research on social learning investigates cases of observational learning, in which subjects are unable to access the food resource during the demonstration phase. Somewhat atypically, our fish could access the food resources during the demonstration phase of the test, mimicking usual foraging conditions in the wild. Thus, shoaling or avoidance could have mediated the discovery of a food location, and a learned association between the food reward and its location would lead to fish subsequently favouring this location [64–66]. The mechanisms underlying different social learning processes are an open question [67,68]. However, from a functional viewpoint, the social learning we describe here and observational learning have the same outcome: both result in individuals’ foraging choices being biased depending on the choices of other individuals. We note that fish without a demonstration, our “control” group, did not form a strong preference for one feeder over the other through repeated feeding, suggesting that demonstrators may have not only biased learning to a particular location but also facilitated learning of that location.

We found extensive population variation in the response to social cues. Depending on the population, social demonstration resulted in copying, avoidance, or no detectable effect on behaviour. Further work is needed to establish the relative contributions of evolution and development to the differences we observed, the underlying neurobehavioural mechanisms, and the question of whether differences in social information use are a byproduct or adaptive specialization. Furthermore, an open question is whether social information use and social learning will vary in parallel: plausibly, in rapidly-changing environments, it may be beneficial to forage with others but not to learn a foraging patch preference from this experience. The differences we observe could have sizable impacts on community dynamics, by shaping and maintaining population-specific foraging preferences or avoidances [69,70]. Our findings also suggest that social learning researchers should pay close attention to the origin and developmental history of their study subjects.

Our dataset is available in the ESM.

## Authors contribution

LCT and SMR conceived and designed the study. LCT carried field work, collected and analysed data from videos, and wrote the first draft of the manuscript. LCT and SMR revised the manuscript. All authors approved submission of the final version.

## Ethical note

We minimized handling time and released fish immediately after testing at their capture location. All procedures were carried out in accordance with Trinidadian and Canadian law, with Animal Behavior Society and Canadian Council for Animal Care guidelines, and were approved by the Animal Care Committee of McGill University (Protocol # 2012-7133).

## Acknowledgements

We thank Maria Cabrera, Lisa Jacquin, Ioannis Leris, Felipe Perez-Jvostov, Dominic Richard, and Paul Q. Sims for help with data collection, Hammond Noriega, Carl and Kelly Fitzjames for field advice, Lauren Redies and Sofija Bekarovska for assistance with video coding, Pierre-Olivier Montiglio for help with data analysis, and Adam Reddon, Will Swaney, Sarah Turner, and Ivon Vassileva for feedback on the experiment and paper.

## References

1. Dall SRX, Giraldeau L-A, Olsson O, McNamara JM, Stephens DW. 2005 Information and its use by animals in evolutionary ecology. Trends Ecol. Evol. 20, 187–93. (doi:10.1016/j.tree.2005.01.010)

2. Hoppitt W, Laland KN. 2013 Social learning: An introduction to mechanisms, methods and models. Princeton: Princeton University Press.

3. Laland KN. 2004 Social learning strategies. Learn. Behav. 32, 4–14.

4. Kendal RL, Coolen I, van Bergen Y, Laland KN. 2005 Trade-offs in the adaptive use of social and asocial learning. Adv. Study Behav. 35, 333–379. (doi:10.1016/S0065-3454(05)35008-X)

5. Rendell L, Fogarty L, Hoppitt WJE, Morgan TJH, Webster MM, Laland KN. 2011 Cognitive culture: Theoretical and empirical insights into social learning strategies. Trends Cogn. Sci. 15, 68–76. (doi:10.1016/j.tics.2010.12.002)

6. Boyd R, Richerson PJ. 1985 Culture and the evolutionary process. Chicago: University of Chicago Press.

7. Webster MM, Laland KN. 2008 Social learning strategies and predation risk: minnows copy only when using private information would be costly. Proc. Biol. Sci. 275, 2869–76. (doi:10.1098/rspb.2008.0817)

8. Chapman BB, Ward AJW, Krause J. 2008 Schooling and learning: early social environment predicts social learning ability in the guppy, *Poecilia reticulata*. Anim. Behav. 76, 923–929. (doi:10.1016/j.anbehav.2008.03.022)

9. Toelch U, Bruce MJ, Newson L, Richerson PJ, Reader SM. 2014 Individual consistency and flexibility in human social information use. Proc. R. Soc. B Biol. Sci. 281, 20132864.

10. Rosa P, Nguyen V, Dubois F. 2012 Individual differences in sampling behaviour predict social information use in zebra finches. Behav. Ecol. Sociobiol. 66, 1259–1265. (doi:10.1007/s00265-012-1379-3)

11. Reader SM. 2015 Causes of individual differences in animal exploration and search. Top. Cogn. Sci. 7, 451–468. (doi:10.1111/tops.12148)

12. Leris I, Reader SM. 2016 Age and early social environment influence guppy social learning propensities. Anim. Behav. 120, 11–19. (doi:10.1016/j.anbehav.2016.07.012)

13. Weigl P, Hanson E. 1980 Observational learning and the feeding behavior of the red squirrel *Tamiasciurus hudsonicus*: The ontogeny of optimization. Ecology 61, 213–8.

14. Laland KN, Williams K. 1998 Social transmission of maladaptive information in the guppy. Behav. Ecol. 9, 493–499. (doi:10.1093/beheco/9.5.493)

15. Giraldeau L-A, Valone TJ, Templeton JJ. 2002 Potential disadvantages of using socially acquired information. Philos. Trans. R. Soc. B Biol. Sci. 357, 1559–1566. (doi:10.1098/rstb.2002.1065)

16. Feldman MW, Aoki K, Kumm J. 1996 Individual versus social learning: evolutionary analysis in a fluctuating environment. Anthropol. Sci. 104, 209–232.

17. Seppanen JT, Forsman JT, Mönkkönen M, Thomson RL. 2007 Social information use is a process across time, space, and ecology, reaching heterospecifics. Ecology 88, 1622–1633.

18. Bouchard J, Goodyer W, Lefebvre L. 2007 Social learning and innovation are positively correlated in pigeons (*Columba livia*). Anim. Cogn. 10, 259–266. (doi:10.1007/s10071-006-0064-1)

19. Reader S. 2003 Innovation and social learning: individual variation and brain evolution. Anim. Biol. 53, 147–158. (doi:10.1163/157075603769700340)

20. Dawson EH, Avarguès-Weber A, Chittka L, Leadbeater E. 2013 Learning by observation emerges from simple associations in an insect model. Curr. Biol. 23, 727–730. (doi:10.1016/j.cub.2013.03.035)

21. Lindeyer CM, Meaney MJ, Reader SM. 2013 Early maternal care predicts reliance on social learning about food in adult rats. Dev. Psychobiol. 55, 168–75. (doi:10.1002/dev.21009)

22. Melo AI, Lovic V, Gonzalez A, Madden M, Sinopoli K, Fleming AS. 2006 Maternal and littermate deprivation disrupts maternal behavior and social-learning of food preference in adulthood: tactile stimulation, nest odor, and social rearing prevent these effects. Dev. Psychobiol. 48, 209–219. (doi:10.1002/dev)

23. Templeton JJ, Kamil AC, Balda RP. 1999 Sociality and social learning in two species of corvids: The pinyon jay (*Gymnorhinus cyanocephalus*) and the Clark’s nutcracker (*Nucifraga columbiana*). J. Comp. Psychol. 113, 450–455. (doi:10.1037//0735-7036.113.4.450)

24. Reader SM, Hager Y, Laland KN. 2011 The evolution of primate general and cultural intelligence. Philos. Trans. R. Soc. Lond. B Biol. Sci. 366, 1017–27. (doi:10.1098/rstb.2010.0342)

25. Coolen I, van Bergen Y, Day RL, Laland KN. 2003 Species difference in adaptive use of public information in sticklebacks. Proc. Biol. Sci. 270, 2413–9. (doi:10.1098/rspb.2003.2525)

26. Webster MM, Chouinard-Thuly L, Herczeg G, Riley R, Rogers S, Shapiro MD, Shikano T, Laland KN. 2019 A four-questions perspective on public information use in sticklebacks (Gasterosteidae). Roy. Soc. Open Sci. 6, 181735 (doi:10.1098/rsos.181735)

27. Marchetti C, Drent PJ. 2000 Individual differences in the use of social information in foraging by captive great tits. Anim. Behav. 60, 131–140. (doi:10.1006/anbe.2000.1443)

28. Trompf L, Brown C. 2014 Personality affects learning and trade-offs between private and social information in guppies, *Poecilia reticulata*. Anim. Behav. 88, 99–106. (doi:10.1016/j.anbehav.2013.11.022)

29. Carlier P, Lefebvre L. 1997 Ecological differences in social learning between adjacent, mixing, populations of Zenaida doves. Ethology 103, 772–784. (doi:10.1111/j.1439-0310.1997.tb00185.x)

30. Dolman CS, Templeton JJ, Lefebvre L. 1996 Mode of foraging competition is related to tutor preference in Zenaida aurita. J. Comp. Psychol. 110, 45–54.

31. Deacon AE, Ramnarine IW, Magurran AE. 2011 How reproductive ecology contributes to the spread of a globally invasive fish. PLoS One 6, e24416. (doi:10.1371/journal.pone.0024416)

32. Camacho-Cervantes M, Ojanguren AF, Magurran AE. 2015 Exploratory behaviour and transmission of information between the invasive guppy and native Mexican topminnows. Anim. Behav. 106, 115–120. (doi:10.1016/j.anbehav.2015.05.012)

33. Reader SM, Kendal JR, Laland KN. 2003 Social learning of foraging sites and escape routes in wild Trinidadian guppies. Anim. Behav. 66, 729–739. (doi:10.1006/anbe.2003.2252)

34. Reznick DN, Rodd FH, Cardenas M, The S, Naturalist A, Mar N, Rodd H. 1996 Life-history evolution in guppies (*Poecilia reticulata : Poeciliidae*). IV. Parallelism in life-history phenotypes. Am. Nat. 147, 319–338.

35. Endler JA. 1984 Natural and sexual selection on colour patterns in poeciliid fishes. Environ. Biol. Fishes 9, 173–190.

36. Torres Dowdall J, Handelsman CA, Ruell EW, Auer SK, Reznick DN, Ghalambor CK. 2012 Fine-scale local adaptation in life histories along a continuous environmental gradient in Trinidadian guppies. Funct. Ecol. 26, 616–627. (doi:10.1111/j.1365-2435.2012.01980.x)

37. Magurran AE. 2005 Evolutionary ecology: the Trinidadian guppy. Oxford: Oxford University Press.

38. Fraser DF, Gilliam JF. 1987 Feeding under predation hazard: response of the guppy and Hart’s rivulus from sites with contrasting predation hazard. Behav. Ecol. Sociobiol. 21, 203–209. (doi:10.1007/BF00292500)

39. Magurran AE, Seghers BH, Shaw PW, Carvalho GR. 1994 Schooling preferences for familiar fish in the guppy, *Poecilia reticulata*. J. Fish Biol. 45, 401–406. (doi:10.1111/j.1095-8649.1994.tb01322.x)

40. Brown GE, MacNaughton CJ, Elvidge CK, Ramnarine I, Godin JGJ. 2009 Provenance and threat-sensitive predator avoidance patterns in wild-caught Trinidadian guppies. Behav. Ecol. Sociobiol. 63, 699–706. (doi:10.1007/s00265-008-0703-4)

41. Magurran, Seghers BH. 1991 Variation in schooling and aggression amongst guppy (*Poecilia reticulata*) populations in Trinidad. Behaviour 118, 214–234.

42. Magurran AE, Seghers BH. 1991 Variation in schooling and aggression amongst guppy (*Poecilia reticulata*) populations in trinidad. Behaviour 118, 214–234.

43. Grether GF, Millie DF, Bryant MJ, Reznick DN, Mayea W. 2001 Rain forest canopy cover, resource availablity, and life history evolution in guppies. Ecology 82, 1546–1559. (doi:10.1890/0012-9658(2001)082[1546:RFCCRA]2.0.CO;2)

44. Jacquin L, Reader SM, Boniface A, Mateluna J, Patalas I, Pérez-Jvostov F, Hendry AP. 2016 Parallel and non-parallel behavioural evolution in response to parasitism and predation in Trinidadian guppies. J. Evol. Biol. 29, 1406–1422. (doi:10.1111/jeb.12880)

45. Gotanda KM, Delaire LC, Raeymaekers JAM, Pe F, Dargent F, Bentzen P, Scott ME, Fussmann GF, Hendry AP. 2013 Adding parasites to the guppy-predation story : insights from field surveys. Oecologia 172, 155–166. (doi:10.1007/s00442-012-2485-7)

46. Willing E, Bentzen P, Oosterhout CVAN. 2010 Genome-wide single nucleotide polymorphisms reveal population history and adaptive divergence in wild guppies. Mol. Ecol. 19, 968–984. (doi:10.1111/j.1365-294X.2010.04528.x)

47. Reznick DN, Rodd FH, Cardenas M. 1996 Life-history evolution in guppies (*Poecilia reticulata: Poeciliidae*): IV. Parallelism in life-history phenotypes. Am. Nat. 147, 319–338.

48. Rodd FH, Reznick DN. 1991 Life-History Evolution in Guppies .3. the Impact of Prawn Predation on Guppy Life Histories. Oikos 62, 13–19. (doi:10.2307/3545440)

49. Reader SM, Biro D. 2010 Experimental identification of social learning in wild animals. Learn. Behav. 38, 265–83. (doi:10.3758/LB.38.3.265)

50. Depasquale C, Neuberger T, Hirrlinger AM, Braithwaite VA. 2016 The influence of complex and threatening environments in early life on brain size and behaviour. Proc. R. Soc. B Biol. Sci. 283, 20152564. (doi:10.1098/rspb.2015.2564)

51. Dussault GV, Kramer D. 1981 Food and feeding behavior of the guppy, *Poecilia reticulata* (Pisces: Poeciliidae). Can. J. Zool. 59, 684–701.

52. R Core Team. 2015 R: A language and environment for statistical computing.

53. Wickham H. 2009 ggplot2: Elegant Graphics for Data Analysis.

54. Bates D, Maechler M, Bolker B, Walker S. 2015 Fitting Linear Mixed-Effects Models Using lme4. J. Stat. Softw. 67, 1–48.

55. Harrison XA. 2015 A comparison of observation-level random effect and Beta-Binomial models for modelling overdispersion in Binomial data in ecology & evolution. PeerJ 3, e1114. (doi:10.7717/peerj.1114)

56. Seppanen JT, Forsman JT, Mönkkönen M, Thomson RL. 2007 Social information use is a process across time, space, and ecology, reaching heterospecifics. Ecology 88, 1622–1633.

57. Boogert NJ, Zimmer C, Spencer KA. 2013 Pre- and post-natal stress have opposing effects on social information use. Biol. Lett. 9, 20121088.

58. Seppanen J-T, Forsman JT, Monkkonen M, Krams I, Salmi T. 2011 New behavioural trait adopted or rejected by observing heterospecific tutor fitness. Proc. R. Soc. B Biol. Sci. 278, 1736–1741. (doi:10.1098/rspb.2010.1610)

59. Durisko Z, Dukas R. 2013 Attraction to and learning from social cues in fruitfly larvae. Proc. R. Soc. B Biol. Sci. 280, 20131398.

60. Millar NP, Reznick DN, Kinnison MT, Hendry AP. 2006 Disentangling the selective factors that act on male colour in wild guppies. Oikos 113, 1–12. (doi:10.1111/j.0030-1299.2006.14038.x)

61. Crispo E, Bentzen P, Reznick DN, Kinnison MT, Hendry AP. 2006 The relative influence of natural selection and geography on gene flow in guppies. Mol. Ecol. 15, 49–62. (doi:10.1111/j.1365-294X.2005.02764.x)

62. Jirotkul M. 1999 Population density influences male-male competition in guppies. Anim. Behav. 58, 1169–1175. (doi:10.1006/anbe.1999.1248)

63. Gamble S, Lindholm AK, Endler JA, Brooks R. 2003 Environmental variation and the maintenance of polymorphism: The effect of ambient light spectrum on mating behaviour and sexual selection in guppies. Ecol. Lett. 6, 463–472. (doi:10.1046/j.1461-0248.2003.00449.x)

64. Atton N, Hoppitt W, Webster MM, Galef BG, Laland KN. 2012 Information flow through threespine stickleback networks without social transmission. Proc. R. Soc. B Biol. Sci. 279, 4272–4278. (doi:10.1098/rspb.2012.1462)

65. Heyes C. M. 1994 Social learning in animals: categories and mechanisms. Biol. Rev. 69, 207–231.

66. Laland K, Williams K. 1997 Shoaling generates social learning of foraging information in guppies. Anim. Behav. 53, 1161–1169.

67. Heyes C. 2012 What’s social about social learning? J. Comp. Psychol. 126, 193–202. (doi:10.1037/a0025180)

68. Reader SM. 2016 Animal social learning: associations and adaptations. F1000Research 5, 2120. (doi:10.12688/f1000research.7922.1)

69. Schmidt KA, Dall SRX, van Gils JA. 2010 The ecology of information: An overview on the ecological significance of making informed decisions. Oikos 119, 304–316. (doi:10.1111/j.1600-0706.2009.17573.x)

70. Thorogood R, Kokko H, Mappes J. 2018 Social transmission of avoidance among predators facilitates the spread of novel prey. Nat. Ecol. Evol. 2, 254–261. (doi:10.1038/s41559-017-0418-x)

