## Supplementary material for "Population differences in how wild Trinidadian guppies use social information and socially learn": Table S1

**Supplementary table**

**Table S1:** Comparison against chance expectation (50%). Models were fitted without intercept. Effect of the combined variable of river and population on the proportion of pecks directed at the demonstrated location for the demonstration (left) or test (right) phase. The estimates are presented on the logit scale. The GLMM included also a correction for overdispersion in the random effects. Significant p-values ($P<0.05$) are presented in bold.

| | **Demonstration phase** | | | | **Test phase** | | | |
|---|---|---|---|---|---|---|---|---|
| | Estimate | Std. Error | z value | P-value | Estimate | Std. Error | z value | P-value |
| Upper Aripo | -2.43 | 1.24 | 1.96 | **0.050** | -2.35 | 1.01 | 2.32 | **0.020** |
| Paria | -1.89 | 1.54 | 1.23 | 0.22 | -0.33 | 1.1636 | 0.28 | 0.78 |
| Lower Aripo | 3.64 | 1.70 | 2.14 | **0.033** | 1.81 | 1.2330 | 1.47 | 0.14 |
| Lower Marianne | -0.77 | 1.05 | 0.73 | 0.46 | 1.06 | 0.8444 | 1.26 | 0.21 |
| Upper Marianne | 0.31 | 1.07 | 0.29 | 0.77 | 0.46 | 0.8640 | 0.53 | 0.60 |
